# Uttroside B, a US FDA-designated ‘Orphan drug’, mitigates the development of hepatocellular carcinoma and its pulmonary metastasis via EGFR/ERK-mediated inhibition of SREBP-1 and STAT-3

**DOI:** 10.1101/2025.07.22.666080

**Authors:** Chenicheri Kizhakkeveettil Keerthana, Tennyson P Rayginia, Kalishwaralal Kalimuthu, Sandhini Saha, Nair H Haritha, Nisthul A Amrutha, Mundanattu Swetha, Sreekumar U Aiswarya, Lekshmi R Nath, Sanjay Suresh Varma, Arun Viswanathan, S Jannet, J Shirley, JS Aparna, Archana Praveen, Vishnu Sunil Jaikumar, Sankar Sundaram, Nikhil Ponnoor Anto, Tushar K Maiti, C Sadasivan, Noah Isakov, Ravi S. Lankalapalli, Kuzhuvelil B Harikumar, Ruby John Anto

**Author notes:** **Corresponding author:** Dr. Ruby John Anto;, **Address for Correspondence:** Dr. Ruby John Anto, Chief Scientist, Centre of Excellence in Nutraceuticals, KSCSTE, Government of Kerala, Bio 360 Life Sciences Park, Thonnakkal, Thiruvananthapuram 695317, Kerala, India.

## Abstract

Hepatocellular carcinoma (HCC) is a highly aggressive tumor with rapid propensity for extrahepatic metastasis, which critically limits the long-term clinical benefits of conventional chemotherapeutics and decreases the overall survival rate of patients. Our previous findings on the exceptional anti-HCC potential and pharmacological safety of uttroside B (Utt-B) have gained multiple international patents and the compound has been designated as an ‘Orphan Drug’ against HCC by the US FDA. The current study substantiates the pharmacodynamics of Utt-B and is the first report to date on the anti-metastatic potential of the compound against HCC. Herein, we demonstrate the role of EGFR/ERK signaling axis and their downstream targets SREBP-1 and STAT-3, the key regulators of HCC development and the pulmonary metastasis, respectively, in orchestrating the anti-HCC and anti-metastatic potential of Utt-B. This is evidenced by the abrogation of the cytotoxic and pro-apoptotic effects of Utt-B upon pharmacological inhibition of this signaling axis. Orthotopic xenograft studies validated that Utt-B treatment restricted the development of tumors via the down-regulation of EGFR/ERK axis. Notably, Utt-B diminishes the migratory and invasive properties of liver cancer cells *in vitro* and impedes the pulmonary metastasis of HCC, *in vivo*. Taken together, the current findings attest to the exceptional therapeutic potential of Utt-B against primary and metastatic HCC and highlight its potential as a candidate drug to be evaluated in the clinics for the benefit of HCC patients having limited prognosis and therapeutic options.

## Introduction

Hepatocellular carcinoma (HCC) accounts for about 90% of malignant liver cancer and is the third leading cause of cancer-related mortality, worldwide [1, 2]. In spite of the significant progress in the therapeutic options for HCC over the past decade, the clinical outcome in terms of overall survival rates of patients with advanced HCC is still unsatisfactory, owing to its high recurrence rates [3] and high propensity of the tumors to directly invade the portal and hepatic veins. Pulmonary metastasis constitutes around 49% of extrahepatic hematogenous metastasis in HCC patients [4, 5]. The prognosis and treatment outcomes in advanced HCC cases with pulmonary metastasis are poor. EGFR over-expression has been reported in over 60% of primary HCC cases [6]. Major survival signaling cascades such as the Ras/Raf/MEK/ERK, JAK/STAT and the PI3K/Akt/mTOR pathway occur downstream of activated EGFR signaling [7]. Interestingly, ERK, a down-stream target of EGFR, is known to activate SREBP-1, the key transcription factor governing various rate-limiting enzymes involved in lipid metabolism [8, 9]. Previous literature suggests that high expression of SREBP-1 leads to excessive *de novo* lipogenesis and lipid uptake which in turn drive the initiation, malignant transformation and progression of HCC [10]. Notably, STAT3, non-canonically activated by receptor tyrosine kinases such as EGFR, plays a pivotal role in epithelial-to-mesenchymal transition and metastatic progression of HCC [11, 12].

Our team has previously isolated and characterized uttroside B (Utt-B) from *S. nigrum* [13]. Notably, our findings on the exceptional therapeutic efficacy of Utt-B against HCC have been granted patents from the USA (US20190160088), Europe (EP3463382), Canada (3,026,426), South Korea (KR1020190008323) and Japan (JP2019520425) [14, 15]. Previous reports by our team suggest that the compound is pharmacologically safe and exhibits superior anti-cancer properties against HCC compared to sorafenib, the US FDA-approved first-line anti-HCC drug [13, 15].

The present study aims to elucidate the pharmacodynamics of Utt-B and evaluate its anti-metastatic potential in the context of HCC, using *in vitro* and *in vivo* pre-clinical models.

## Materials and Methods

### Chemicals

Isolation and purification of Utt-B was done as previously reported [13]. Cell culture reagents such as Dulbecco’s Modified Eagle Medium (DMEM) (GIBCO, 12800-017), Minimum Essential Medium (MEM) (61100061), Fetal Bovine Serum (GIBCO 10270-106), streptomycin sulfate (GIBCO, 11860-038), were obtained from Invitrogen Corporation (Grand Island, USA). Poly Excel HRP/DAB detection system universal kit (PathnSitu Biotechnologies Pvt. Ltd, India, OSH001) was used for immunohistochemistry experiments. MTT reagent (D0801) was purchased from TCI Chemicals (India) Pvt. Ltd. Matrigel (356231) was procured from Corning. Antibodies against β-actin (12620S), p-EGFR-Tyr1068 (3777S), EGFR (2239S), p-ERK1/2-Thr202/Tyr204 (9101S), ERK1/2 (9102S), p-mTOR-Ser2481 (2974S), and p-STAT3-Ser727 (9136S) were obtained from Cell Signaling Technologies (Beverly, MA, USA) and the antibodies against PCNA (sc25280), Ki67 (sc23900), p-EGFR-Tyr1045 (sc57541), STAT3 (sc8019), SREBP-1 (SC-13551), Cyclin D1 (sc8396), E-cadherin (sc8426), MMP2 (sc13594), and MMP9 (sc6840), and Annexin V-FITC/PI apoptosis detection kit (sc4252AK) were purchased from Santa Cruz Biotechnology (Santa Cruz, CA, USA). Antibody against vimentin (MA3 41 745) was purchased from Invitrogen. FITC and Rhodamine conjugated anti-mouse and anti-rabbit secondary antibodies were procured from Calbiochem. DeadEnd™ Fluorimetric TUNEL System was procured from Promega (G3250). MycoSensor qPCR Assay Kit (302107) used to test mycoplasma contamination in cells was purchased from Agilent Technologies. Power SYBR Green PCR master mix (4367659) and High-capacity cDNA kit (4368814) were purchased from Applied Biosystems. U0126 (662005) was purchased from Merck Millipore. All other chemicals were purchased from Sigma Chemicals (St. Louis, MO, USA) unless mentioned otherwise.

### Cell Culture

The liver cancer cell lines, HepG2, Huh7 and Hep3B were from obtained from NCCS, Pune, India. All the cell lines were cultured in Dulbecco’s Modified Eagle Medium (DMEM) supplemented with 10% Fetal Bovine serum. All the cell lines used in this study were mycoplasma-tested and were free from mycoplasma contamination.

### Animal experiments

*In vivo* experiments were conducted in accordance with institutional animal ethical committee guidelines (IAEC No: IAEC/765/RUBY/2019 and IAEC/944/HARI/2023). Mice were housed in a 12 h light/dark cycle with access to standard food pellets and autoclaved water *ad libitum*. 4-5 weeks old (body weight around 24-28g) NOD-SCID Gamma (NOD.CB17-Prkdcscid/J Gamma) male mice were used for the orthotopic model of HCC experiment. The mice were anaesthetized using ketamine (100 mg/kg body weight) and xylazine (16 mg/kg body weight). An incision was made in the abdominal region. Xenografts were established by injecting 2×10^6^ Huh 7 cells (dissolved in 1:1 PBS: Matrigel) into the left hepatic lobe using a Hamilton syringe. The incision was closed using surgical clips. The mice were monitored until they regained consciousness. The surgical clips were removed after the wound was completely healed after 5-7 days post-surgery. One-week post-cell injection, the mice were randomly separated into 2 groups (n=6). Group I was treated with PBS alone, Group II received an intraperitoneal injection (IP) of Utt-B dissolved in PBS at a dose of 10 mg/Kg body weight on alternate days. Drug treatment was continued for up to 4 consecutive weeks, after which animals were euthanized and the blood and tissue samples were collected for further analyses. Murine model of HCC metastasis was generated by injecting 1 × 10^6^ Hep3B cells (dissolved in 1:1 PBS: Matrigel) via the lateral tail vein of 4-5 weeks old NSG male mice. Two days i.e., 48h post-injection, the mice were separated into 2 groups (n=6). Group I received the vehicle, and Group II received an intraperitoneal injection of Utt-B in PBS (10 mg/kg body weight) respectively, on alternate days for 2 months, followed by the euthanasia of animals and subsequent procurement of the tissue samples for further analyses.

### MTT assay

MTT assay to measure the cell viability, and the extent of cytotoxicity induced by different compounds was performed as previously described [14]. The sub-toxic concentration of each inhibitor required to inhibit the function of the respective target molecule (1μM Erlotinib, 5nM rapamycin, and 2μM U0126) was chosen based on the pre-existing literature. The HepG2 cells were pre-treated with the respective inhibitor 6h prior to the treatment with 500nM Utt-B. The cells were incubated for 72h post-treatment with Utt-B.

### Detection of apoptosis by Annexin V-FITC/PI fluorescence microscopy

The presence of apoptotic cells was detected with the help of Annexin V-FITC/PI staining followed by fluorescent microscopy using Annexin V-FITC/PI kit according to the manufacturer’s protocol (Santa Cruz, CA, USA). Curcumin at a dose of 25μM was used as the positive control for the apoptosis assay using Annexin V/PI staining.

### Immunofluorescence studies

Immunofluorescence was used to assess the differential expression of the proteins of our interest in response to Utt-B treatment was performed as previously described [14, 16].

### Wound healing assay

The wound healing assay was performed as previously described [14].

### Boyden chamber cell migration and invasion assay

Migration and invasion assays were performed in 24-well plate with Transwell inserts (8 μm pore size) as previously described [17]. The upper side of the insert was coated with 40μl of Matrigel (Corning) at 37 °C for 30 min for the invasion assay. Matrigel was diluted using serum free DMEM and coated to the inserts at a final concentration of 1mg/ml.

### Cell adhesion assay

Cell adhesion assay was performed using 96-well plates were coated with Matrigel as previously described [17].

### RNA Isolation and cDNA synthesis

RNA isolation from cells was performed by TriZol method. cDNA synthesis was done using High-Capacity cDNA synthesis kit (Applied Biosystems).

### Real-Time PCR

Real-Time PCR was performed using Power SYBR Green PCR master mix, Applied Biosystems (Cat. No. 4367659). The housekeeping gene, GAPDH, was used as the reference gene for normalization. The data were calculated using the 2^−ΔΔ^C_t_ method [18]. The details of the primers used in the study have been included in **Supplementary Table 1**.

### Nanostring nCounter Assay

For NanoString analysis, samples were quantified using Qubit RNA HS assay (Invitrogen, Cat # Q32855) kit and qualitatively analysed on Agilent 2100 bioanalyzer nano chip (Agilent, Cat # 5067-1511). Codeset reaction for nCounter Custom Gene Panel (NanoString Technologies) was carried out as per the Manual (nCounter Gene Expression Panel and custom codeset user manual, MAN-10056-06). RNA was hybridized with Reporter and Capture probes at 65^0^C for 18 hours and samples run on nanoString nCounter SPRINT machine through service provider Theracues Innovations Pvt Ltd, Bangalore, India. The Raw data files were downloaded from nCounter SPRINT machine and analyzed using nSolver™ 4.0 (NanoString Technologies) software. The fold change values and unpaired Student’s t test-based p-value were calculated using the ‘calculate ratio’ module from nSolverTM 4.0 Analysis Software (MAN C0019-08). Differentially expressed genes were identified with thresholds of p-value 1.2 (upregulated) or Fold change negative probe counts. All statistical analysis was performed by using R (version 4.0.2, https://www.r-project.org/). P-value less than 0.05 was considered statistically significant.

### Phosphoproteomics analysis

Phosphoproteomic analysis was carried out using metal oxide affinity enrichment coupled with LC-MS/MS as previously described [19].

### Western Blotting

Western blotting analyses were carried out using specific antibodies as previously described [14, 20].

### Measurement of haematological and serum biochemical parameters

Analysis of the liver function, and renal function profiles of the mice were performed using Animal Biochemistry Analyzer (DryChem NX-500; Fuji Film).

### H&E staining

Haematoxylin and Eosin staining was performed as previously described [14]. Pathological verification of tumor and tissue samples for all the animal experiments were sent for a blinded analysis to the pathologist.

### Sirius Red staining

Sirius Red staining was performed for detecting collagen deposition and fibrotic changes in tissue sections was performed as previously described [21].

### TUNEL assay

TUNEL assay was performed to detect apoptosis in formalin fixed, paraffin-embedded liver tissue sections using Dead End Fluorimetric TUNEL System (Promega) following the manufacturer’s instructions.

### Immunohistochemistry

Immunohistochemical analysis of proteins in the tissue sections was done using the PolyExcel HRP/DAB detection system universal kit for mouse primary antibodies (PathnSitu Biotechnologies Pvt. Ltd, India) according to the manufacturer’s protocol.

### *In silico* docking studies

*In silico* docking studies were performed using the program Schrodinger suite (Maestro 10.4). The crystallographic structures were retrieved from Protein Data Bank for the study. The structures were corrected using protein preparation wizard which ensures the structural accuracy of the initial protein structure. Crystallographic waters were removed, missing hydrogens were added and minimization was performed using OPLS3 forcefield. Chemical structure of Utt-B (CID:44566638) was downloaded from PubChem and prepared using the Ligprep module. Receptor grid was generated around the co-crystallized ligand with the receptor grid generation module. Molecular docking studies were performed using Glide in standard precision (SP) mode.

### Statistical analysis

Data represents the results for experiments performed in triplicate. The quantification of Western blots and immunohistochemistry images were carried out using Image J software. The statistical analysis was performed using Graph Pad Prism software (Graph Pad Software Inc., San Diego, CA, USA). Statistical tests used were Student’s t test, One way ANOVA, Two-way ANOVA etc. Statistical significance was defined as ****P-values≤0.0001, ***P-values≤0.001, **P-values≤0.01 and *P-values ≤0.05; and ns represents non-significance. The error bars represent ± SD, taken from three independent experiments.

## Results

### Uttroside B exhibits anti-HCC effects by targeting EGFR, mTOR and MAPK pathways

Utt-B-mediated changes at the transcriptome level were assessed by performing a Nanostring nCounter assay using HepG2 cells treated with the IC50 concentration of the compound (500nM). The results of the nCounter assay revealed that Utt-B significantly lowers the expression of several genes associated with HCC progression, including, EGFR, BRAF, TGF-β1, JAK1, STAT3, VEGF, JUN, MYC, HIF-1α, CTNNB1 and FOXA1. Interestingly, Utt-B treatment also induced a marked decrease in the expression of CDH2, SNAI1 and MMP3, the most prominent molecular markers associated with cancer metastasis. Further, the pro-apoptotic gene, BAD and anti-fibrotic gene, LEFTY2 were also up-regulated upon Utt-B treatment, suggesting the pro-apoptotic potential of the compound. Additionally, several other genes associated with the poor prognosis of HCC were also found to be down-regulated upon treatment with Utt-B **(Supp. Fig. 1A)**. The functional analysis and STRING analysis of the Nanostring dataset using KEGG and Reactome pathway databases revealed that EGFR, MAPK and JAK/STAT pathways are the major pathways being down-regulated in response to Utt-B **(Fig. 1A-B).** The phosphoproteomic profiling of the control and Utt-B-treated HepG2 cells revealed that a total of 192 unique phosphoproteins were expressed in the control cells, and 211 phosphoproteins were expressed in cells treated with Utt-B, while a total of 584 phosphoproteins were commonly expressed in both groups **(Fig. 1C; Supp. Fig. 1B-C).** Further, a kinome extraction from the phosphoproteomics dataset was performed to delineate Utt-B-mediated changes occurring in the phosphorylation signaling networks of the major protein kinases. The major kinases enriched in the control condition include EGFR, MAPK and mTOR. The expression of PAK1, an inducer of the MAPK survival signaling cascade [22], was observed in the control condition. On the contrary, there was paradigm shift in the enriched kinases in the cells treated with Utt-B. EGFR, mTOR and major MAPK kinases were absent in the treated condition. Most of the enriched kinases in the Utt B-treated group were pro-apoptotic kinases. Interestingly, the expression of TAK1, a prominent kinase involved in the transient phosphorylation and endocytosis of the EGFR [23], was observed in the Utt-B treated samples. Moreover, neither EGFR, nor mTOR and MAPK were present in the treated condition **(Fig. 1D).** Thus, the results of the phosphoproteomics analysis suggest that the major molecular targets of Utt-B are, EGFR, mTOR and MAPK. Immunocytochemical analysis revealed that Utt-B induced a time-dependent decrease in the expression of p-EGFR (Y1045), p-mTOR (S2481) and p-ERK1/2 (T202/Y204) in HepG2 cells **(Fig. 2A-C).** *In silico* molecular docking analyses were performed to decipher the possible direct physical interaction of the compound with any of its molecular targets. As Utt-B is a bulky molecule with a molecular weight of 1215.34 Da, it was observed that the compound was much larger than the active sites of ERK1/2 and STAT3 and hence, the docking was not feasible. However, when the compound was docked with EGFR, it was observed that Utt-B could bind in the extracellular domain of EGFR at the same site, where cetuximab, the anti-EGFR monoclonal antibody binds **(Fig. 2D).** The glide gscore value of Utt-B with EGFR is −7.41 kcal/mol. Interestingly, Utt-B also exhibited binding feasibility in the kinase domain of EGFR, at the same site where erlotinib, the pharmacological inhibitor of EGFR binds **(Fig. 2E).** The glide gscore value of Utt-B is −9.77 kcal/mol, which is slightly more negative than the gscore of erlotinib (−9.25 kcal/mol), indicating better binding affinity and stability of interaction between Utt-B and EGFR, over erlotinib. Further, the docking studies of Utt-B with mTOR revealed that Utt-B could bind at the same inhibitory site in the kinase domain of mTOR, where the pharmacological inhibitor of mTOR, rapamycin binds **(Fig. 2F).** The glide gscore value of Utt-B is −8.05 kcal/mol, which is far greater than that of rapamycin (−15.9 kcal/mol), indicating the better binding affinity and stability of interaction between rapamycin and mTOR, over Utt-B. Hence, the *in silico* findings suggest the possibility of direct physical interaction of Utt-B with EGFR rather than mTOR, owing to the better docking fitness.

**Figure 1:**
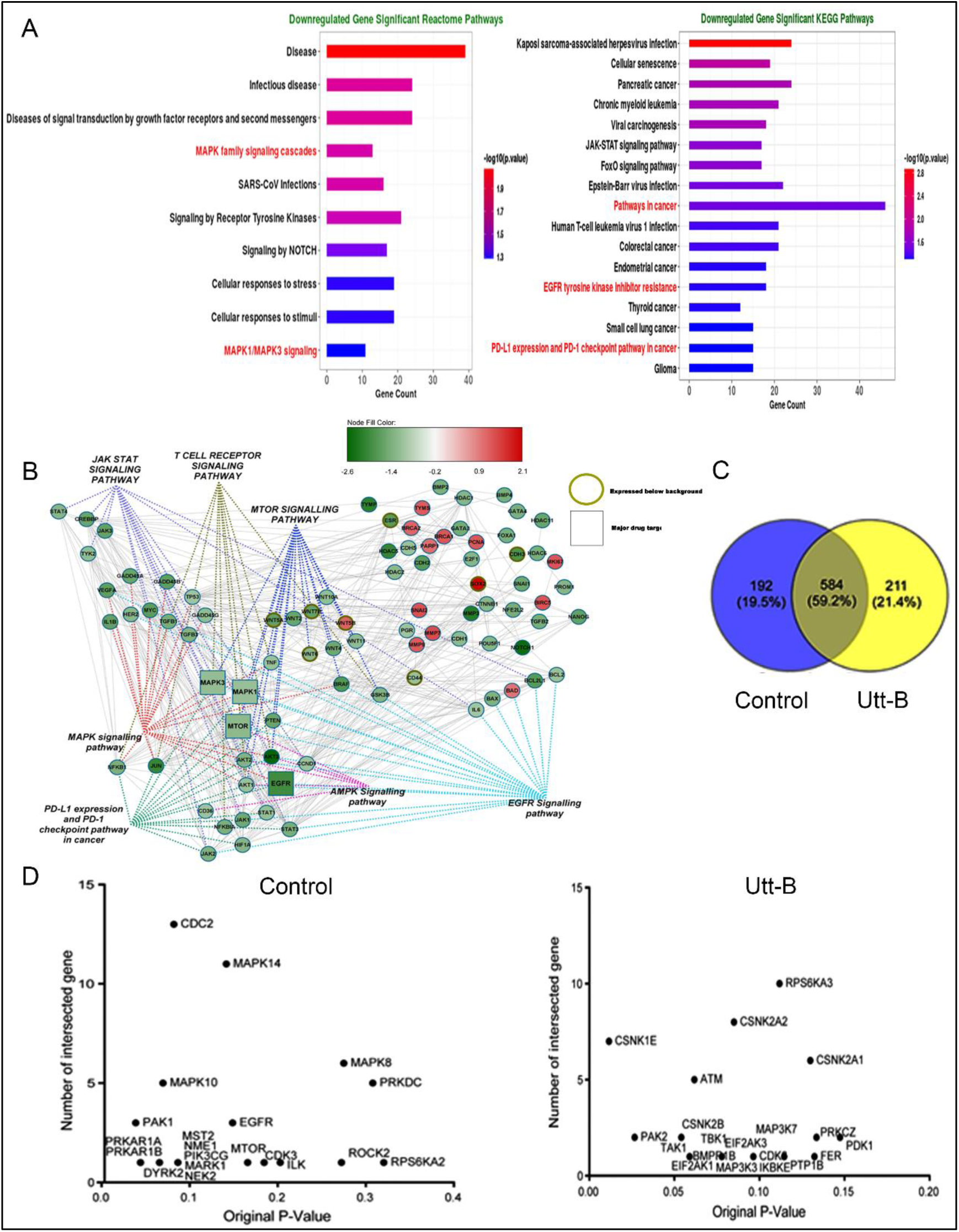
Trancriptomic and phosphopreoteomic analyses reveal that EGFR, MAPK and mTOR pathways are the major targets of Utt-B. **(A)** Pathway analysis of Nanostring nCounter assay data using KEGG and Reactome databases depicts the list of significantly down-regulated pathways in HepG2 cells treated with Utt-B, compared to control. **(B)** STRING analysis showing direct interactions of the down-regulated genes with major signaling pathways associated with the progression of HCC and induction of anti-tumor immune response. **(C)** Phosphoproteomics analysis depicting the number of differentially expressed phosphoproteins in the control and Utt-B-treated HepG2 cells. **(D)** Kinome extraction data showing the enriched kinases unique to the control and Utt-B treatment groups.

**Figure 2:**
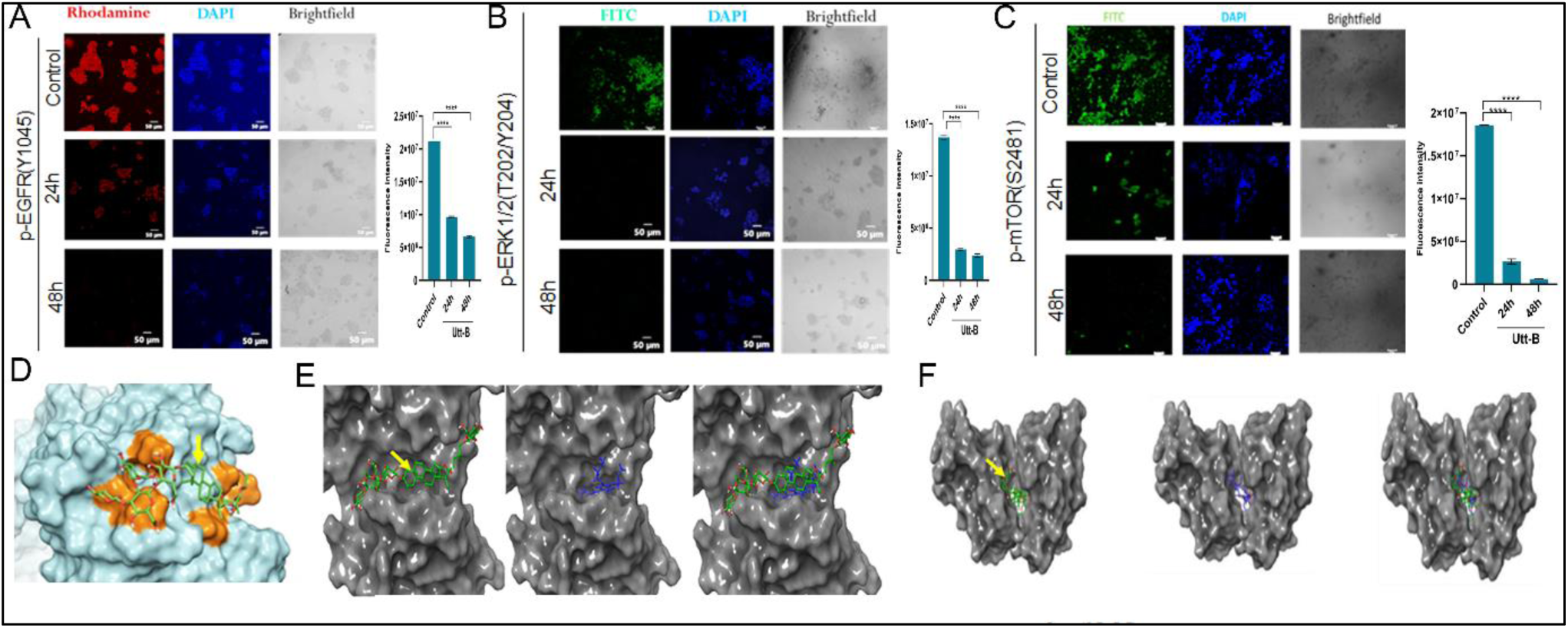
*In vitro* and *in silico* studies indicate the inhibitory effects of Utt-B on EGFR, ERK and mTOR. Immunocytochemical analysis showing the time-dependent changes in the phosphorylation of **(A)** p-EGFR, **(B)** p-ERK1/2 and **(C)** p-mTOR in HepG2 cells. **(D)** Docking pose of Utt-B with the extracellular domain of EGFR as analyzed by in silico studies. Utt-B (indicated in green color and pointed by yellow arrow) binds at the inhibitory site in the extracellular domain of EGFR and forms three H-bonds with Q384, S418 and K465 (indicated in orange color) residues. **(E)** Docking pose and fitness of Utt-B with the kinase domain of EGFR as analyzed by *in silico* studies. From left, docking pose and orientation of Utt-B (indicated in green color and pointed by yellow arrow) binding at the inhibitory site in the kinase domain of EGFR, docking pose and orientation of erlotinib indicated in blue color) with EGFR and comparative docking pose and orientation of erlotinib and Utt-B with EGFR, depicting that Utt-B binds at the same site as that of erlotinib. **(F)** Docking pose and fitness of Utt-B with the kinase domain of mTOR as analyzed by *in silico* studies. From left, docking pose and orientation of Utt-B (indicated in green color and pointed by yellow arrow) binding at the inhibitory site in the kinase domain of mTOR, docking pose and orientation of rapamycin indicated in blue color) with mTOR, and comparative docking pose and orientation of rapamycin and Utt-B with mTOR, depicting that Utt-B binds at the same site as that of rapamycin. One-way ANOVA followed by Tukey’s post hoc t test analysis was used for statistical comparison between different groups. ****P ≤ 0.0001; **P ≤ 0.01; *P ≤ 0.05; ns-non-significant.

### Pharmacological inhibition of EGFR/ERK axis abrogates the anti-HCC potential of uttroside B, attesting the regulatory role of this signaling axis

Administration of Utt-B in the absence of the chemical inhibitors resulted in 50% cytotoxicity and prominent induction of early apoptosis in HepG2 cells. Although, each of the chemical inhibitors induced cytotoxicity and mild apoptosis that was statistically significant, none of the inhibitors decreased the cell viability below 70%, suggesting that the chosen concentrations were suitable for the pharmacological inhibition studies. However, administration of Utt-B to cells pre-treated with 1μM erlotinib (Erlo), drastically decreased the profound cytotoxic and pro-apoptotic activities of Utt-B. The resulting cytotoxicity and apoptosis in the cells treated with the combination of Erlo+Utt-B were comparable to that of cells treated with Erlo alone and was far lesser compared to that of the cells treated with Utt-B alone. These observations confirm that Utt-B-mediated down-regulation of EGFR is responsible for its anti-HCC effects **(Fig. 3A-B).** Based on the earlier findings of the present study, it is evident that both mTOR and MAPK pathways that operate down-stream of EGFR, have a crucial role in the action of Utt-B. Inhibition of the mTOR pathway using rapamycin (5nM Rap) revealed that the cytotoxic and pro-apoptotic effects produced by Rap alone was comparable to Erlo alone, and was far lesser than the effects produced by Utt-B alone in the absence of Rap. Surprisingly, the administration of Utt-B to cells pre-treated with Rap resulted in the enhancement of cytotoxicity and apoptosis. These results clearly demonstrate that mTOR does not play a regulatory role in dictating the anti-HCC potential of Utt-B. However, the down-regulation of mTOR could be a consequence of Utt-B-mediated down-regulation of EGFR, the up-stream regulator of the mTOR pathway **(Fig. 3 C-D).** Further, the MAPK/ERK pathway was inhibited using 2μM U0126. The cytotoxic and pro apoptotic effects produced by U0126 alone was comparable to Erlo alone and Rap alone, and was far lesser than the effects produced by Utt-B alone in the absence of U0126. However, when Utt-B was administered to cells pre-treated with 2μM U0126, the profound cytotoxic and pro-apoptotic activities of Utt-B were diminished, despite the presence of EGFR. The resulting cytotoxicity and apoptosis in the cells treated with the combination of U0126+Utt-B was comparable to that of cells treated with U0126 alone and was far lesser compared to that of the cells treated with Utt-B alone **(Fig. 3 E-F)**. Notably, immunoblotting analyses validated the time-dependent down-regulation of p-EGFR (Y1068,1045) and p-ERK1/2 (T202/Y204) and the concomitant down-regulation of SREBP-1 in response to Utt-B treatment **(Fig. 3 G).** Taken together, the current findings confirm that Utt-B-mediated down-regulation of the EGFR/ERK signaling axis and its down-stream target, SREBP-1 contribute to the remarkable anti-cancer effects elicited by the compound, possibly via limiting excessive *de novo* lipogenesis and aberrant lipid metabolism.

**Figure 3:**
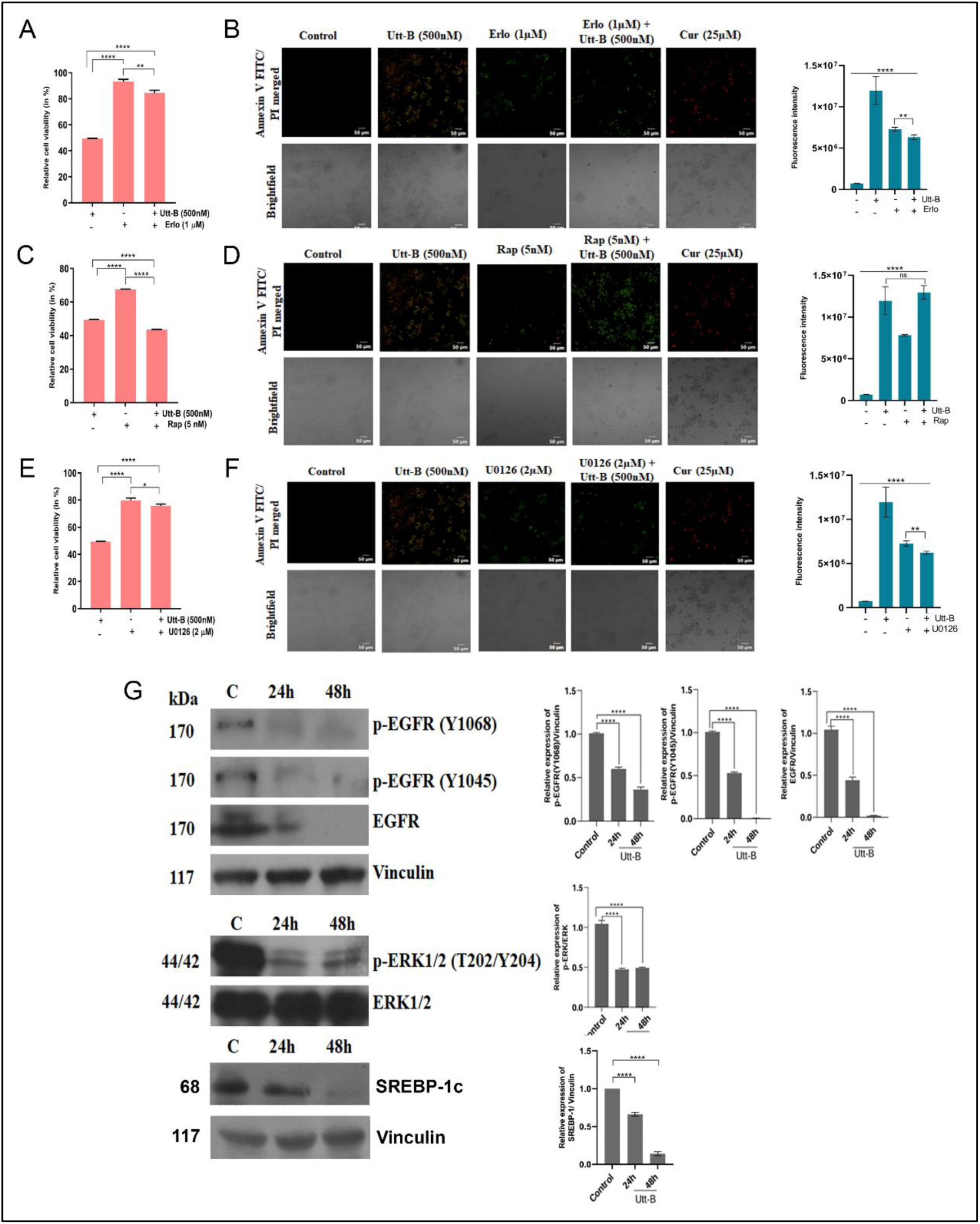
Pharmacological inhibition of EGFR/ERK axis abrogates the therapeutic effects of Utt-B against HCC. MTT assay depicting the changes in Utt-B-induced cytotoxicity in HepG2 cells upon pharmacological inhibition of **(A)** EGFR with erlotinib, **(C)** mTOR with rapamycin and **(E)** MAPK/ERK with U0126. Annexin V/PI staining showing the changes in Utt-B-mediated induction of apoptosis in HepG2 cells upon pharmacological inhibition of **(B)** EGFR with erlotinib, **(D)** mTOR with rapamycin and **(F)** MAPK/ERK with U0126. One-way ANOVA followed by Tukey’s post hoc t test analysis was used for statistical comparison between different groups. ****P ≤ 0.0001; **P ≤ 0.01; *P ≤ 0.05; ns-non-significant. **(G)** Western blot showing the down-regulation of EGFR/ERK axis and SREBP-1 in HepG2 cells in response to Utt-B treatment. Student’s t test was used for statistical comparison between different groups. ****P ≤ 0.0001; ***P ≤ 0.001; *P ≤ 0.05.

### Uttroside B impedes the development of orthotopic hepatic tumors and induces apoptosis*, in vivo*

*In vivo* orthotopic xenograft model of HCC which facilitate a better understanding of the therapeutic effects of drugs in the native tumor microenvironment, was established in NSG mice **(Supp. Fig. 1D)**. Primary solid tumors, which were verified as HCC by histopathological analysis, were observed on the livers of the untreated animals. Interestingly, only one animal out of the six animals, from the Utt-B-treated group developed solid hepatic tumors **(Fig. 4A)**. The tumor sections from this animal were used for all further analyses. The liver: body weight ratio was much lesser in the treatment group, indicating a positive therapeutic response to Utt-B **(Fig. 4B)**. Histopathological analysis of the tumor tissues and the corresponding liver tissues revealed the presence of well differentiated solid hepatic tumor in the control group. Presence of sheets of malignant cells showing mitosis were noted in the hepatic tumor tissues collected from all the control mice and from the only tumor-bearing animal in the Utt-B group. Significant congestion was observed in the corresponding liver tissues from the control group, while it was mild in that of the tumor-bearing animal from Utt-B group. Livers of all other mice in the Utt-B group showed normal histology **(Fig. 4C)**. Excessive collagen deposition associated with the pathogenesis of HCC was noted in the livers of control mice, while it was significantly less in the livers of mice treated with Utt-B, as assessed by Sirius Red staining. This observation strongly indicate that Utt-B improves the histopathological features of the livers of mice bearing orthotopic HCC xenografts **(Fig. 4D)**. The elevated levels of liver enzymes such as, SGOT, SGPT, ALP and increased levels of total bilirubin, creatinine and blood urea nitrogen in the serum serve as markers of impaired hepatic and renal dysfunction. The biochemical analysis of the serum indicates a drastic increase in the levels of the liver enzymes, SGOT, SGPT and the levels of total bilirubin, blood urea nitrogen and creatinine in the control mice, while, these levels were stabilized well-within the normal range in the mice treated with Utt-B **(Fig. 4 E-F; Supp. Fig. 1E-F)**. These results suggest that Utt-B stabilizes hepatic and renal function in mice bearing orthotopic HCC xenografts. Immunohistochemical analysis revealed that the expression of the HCC specific biomarker, alpha-foetoprotein (AFP) and the proliferation markers, Ki-67 and PCNA, were significantly high in the control tumor tissue while the expression of all these proteins were markedly reduced in the tumor tissue collected from the only tumor-bearing animal in the Utt-B treatment group, attesting the remarkable efficacy of Utt-B against HCC **(Fig. 4G)**. Furthermore, a drastic reduction in the expression of p-EGFR and p-ERK1/2 was observed in the tumor tissue collected from the only tumor-bearing animal in the Utt-B group, compared to the control group **(Fig. 4H)**. Moreover, Utt-B induced drastic DNA fragmentation, a classical hallmark of apoptosis, in the liver tissues of mice bearing orthotopic HCC xenografts, as assessed by TUNEL assay **(Fig. 4I)**. While 50% necrosis was observed in the histopathological analysis of the hepatic tumor tissues collected from the control animals, there were no signs of necrosis in the tumor tissue collected from the only tumor-bearing animal in the Utt-B group. This could be due to the extensive apoptosis induced by Utt-B as evidenced by the results of the TUNEL assay.

**Figure 4:**
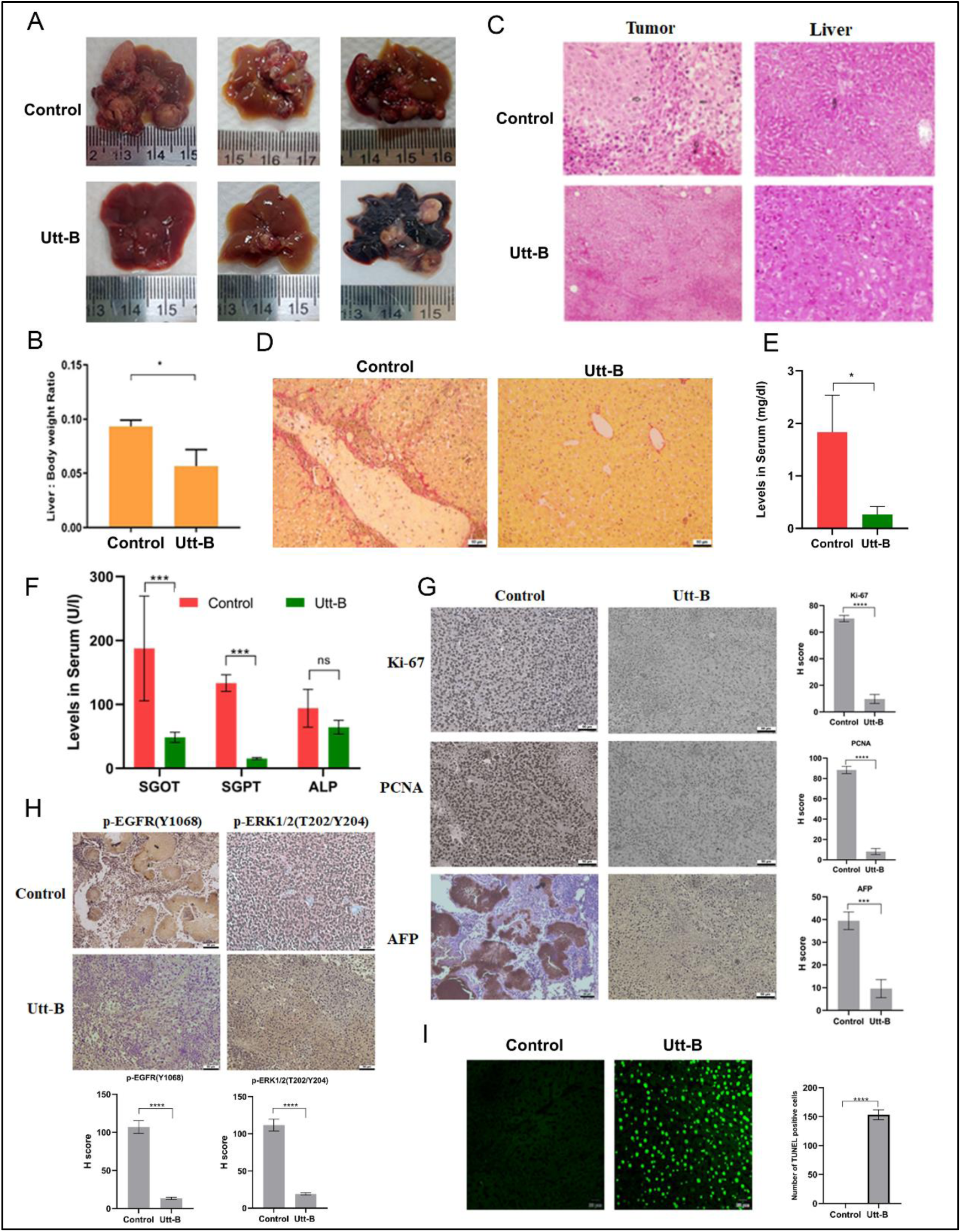
Utt-B hinders the development of hepatic tumors in a murine orthotopic xenograft model of HCC via down-regulation of the EGFR/ERK axis. **(A)** Representative images of livers bearing orthotopic hepatic tumors in control and Utt-B-treated mice from the orthotopic xenograft model. **(B)** Graph depicting changes in the liver: body weight ratio in mice from control and treatment groups. Mann-Whitney test was used for statistical comparison between different groups. *P ≤ 0.05. **(C)** Histopathological analysis of orthotopic tumor and corresponding liver tissues from control group and the tumor collected from the only tumor-bearing animal from Utt-B treated group of the orthotopic xenograft model. **(D)** Sirius Red staining depicting the excessive deposition of collagen in the livers of control mice and its minimal deposition in livers of mice treated with Utt-B. **(E-F)** Levels of bilirubin and liver enzymes from the control and Utt-B treated groups as assessed by biochemical analysis of serum samples. **(G)** Immunohistochemical analysis of cancer biomarkers, Ki-67, PCNA and HCC specific biomarker, AFP in the tumor tissues of control and Utt-B treated mice from the orthotopic xenograft model. **(H)** Immunohistochemical analysis showing the relative tissue expression of p-EGFR and p-ERK1/2 in the tumor tissues of control and Utt-B-treated mice from the orthotopic xenograft model. **(I)** TUNEL assay showing Utt-B-mediated DNA fragmentation and apoptosis in the livers of mice treated with Utt-B. Student’s t test was used for statistical comparison between different groups. ****P ≤ 0.0001; ***P ≤ 0.001; *P ≤ 0.05; ns-non-significant.

### Uttroside B retards the rate of migration, invasion and epithelial-mesenchymal transition (EMT) of HepG2 cells, *in vitro*

Our next attempt was to evaluate the efficacy of Utt-B against invasion and metastasis. The anti-migratory effects of Utt-B was studied using a wound closure assay, where no significant wound closure was observed in the cells treated with 500nM Utt-B, even after 48h post-treatment **(Fig. 5A).** The anti-migratory effects of Utt-B was further confirmed by a cell migration assay, where a drastic reduction in the chemotactic migration was noted in HepG2 cells treated with Utt-B **(Fig. 5B)**. Further, Utt-B led to a marked decrease in the number of cells that penetrate across the ECM barrier and migrate towards the chemoattractant, in the cell invasion assay **(Fig. 5C)**. Interestingly, Utt-B led to a drastic reduction in the number of cells that adhered on to the surface of Matrigel-coated wells in the cell adhesion assay, validating the inhibitory effect of Utt-B on the post-invasion adhesion potential of liver cancer cells, and confirming the anti-metastatic property of Utt-B, *in vitro* **(Fig. 5D)**. The results of the qRT-PCR revealed that Utt-B decreases the mRNA expression levels of mesenchymal markers such as N-cadherin, TWIST2 and SNAI1 **(Fig. 5E)**. The immunoblotting results indicate that Utt-B induces significant up-regulation of the epithelial marker, E-cadherin and down-regulation of the mesenchymal marker, vimentin, in a time-dependent manner **(Fig. 5F)**. The reduction in the expression of mesenchymal markers and enhancement in the expression of epithelial marker suggests the potential of Utt-B in preventing the epithelial-to-mesenchymal transition, one of the major pre-requisites for the initiation of metastasis. Previous literature suggests the phosphorylation of STAT3 at the S727 residue is commonly mediated by ERK1/2 [24]. Utt-B induces a time-dependent down-regulation of p-STAT3 (S727) and its down-stream target, Cyclin D1 **(Fig. 5G)**, suggesting that Utt-B could be preventing the metastasis of HCC, via down-regulation of the EGFR/ERK axis and the subsequent down-regulation of STAT3 pathway. This could be one of the possible mechanisms underlying the strong anti-metastatic potential of Utt-B.

**Figure 5:**
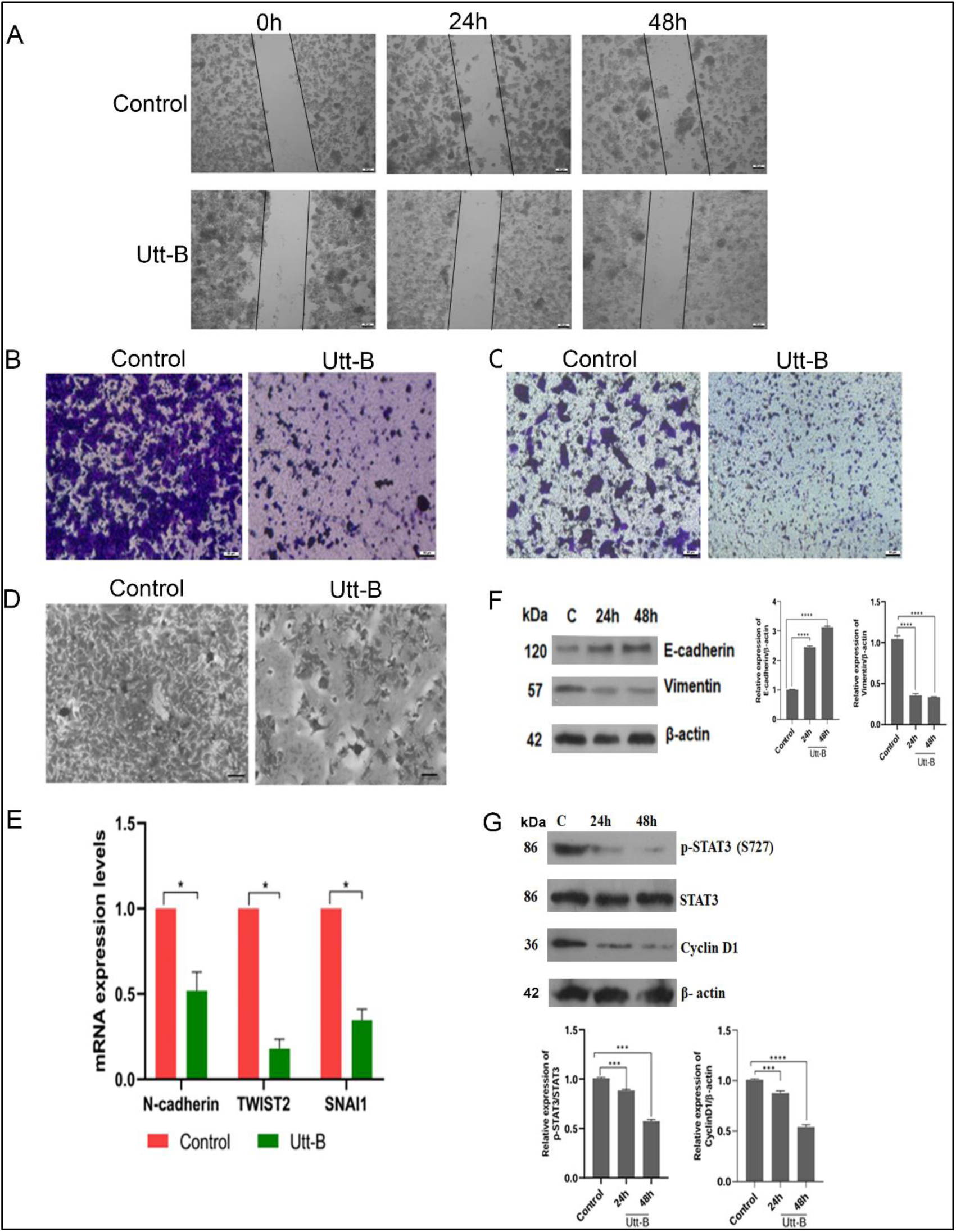
Utt-B significantly inhibits the migration, invasion and adhesion and epithelial-mesenchymal transition of HepG2 cells, via down-regulation of STAT 3 pathway. **(A)** Wound healing assay depicting the ability of Utt-B in inhibiting the cell migration in treated HepG2 cells as compared to the untreated control cells. **(B)** Transwell migration assay indicating the anti-migratory effects of Utt-B in HepG2 cells. **(C)** Transwell invasion assay indicating the anti-invasive properties of Utt-B in HepG2 cells. **(D**) Cell adhesion assay demonstrating the strong inhibitory effects of Utt-B against the adhesion of HepG2 cells onto Matrigel coated wells. **(E)** Relative fold change in the mRNA expression levels of EMT markers in HepG2 cells treated with Utt-B as analyzed using qRT-PCR. **(F)** Western blot analysis showing the Utt-B-mediated changes in the expression of the EMT markers, E-cadherin and Vimentin in HepG2 cells. **(G)** Western blot showing the down-regulation of STAT3 pathway in HepG2 cells in response to Utt-B treatment. Student’s t test was used for statistical comparison between different groups. ****P ≤ 0.0001; ***P ≤ 0.001; *P ≤ 0.05.

### Uttroside B alleviates the pulmonary metastasis of HCC, *in vivo*

An *in vivo* murine metastasis model of HCC was established in NSG mice using the invasive human liver cancer cell line, Hep3B **(Supp. Fig. 1G)**. Solid tumors and large colorless nodules that are indicative of HCC metastasis were developed on the lungs of all untreated animals, while only one animal out of the six animals from the Utt-B group developed secondary solid tumors on the lungs. Moreover, the tumor nodules developed on the lungs of this animal were much lower in size as compared to that of the control mice. The lung tumor sections from this animal were used for all further analyses **(Fig. 6A-B)**. Histopathological analysis of the metastatic tumors and the corresponding lung tissues validated the occurrence of solid tumors in all the mice from the control group. The tumor nodules isolated from the only tumor-bearing animal from the Utt B group were also identified as HCC. The lungs of control mice showed mild collapse, while that of Utt-B-treated mice showed normal histology. Moreover, the control tumors indicated 50% necrosis whereas, no traces of necrotic changes were visible in the tumor tissue of the only tumor-bearing animal from the Utt-B group. Tumor necrosis has a positive correlation with intra-tumor hypoxia, vascular invasion and aggressiveness of tumor metastasis [25]. Therefore, the current results confirm that Utt-B drastically decelerates the development of aggressive metastatic tumors in the lungs, *in vivo* **(Fig. 6C)**. The levels of SGOT, SGPT, ALP and total bilirubin were stabilized within the normal range in mice treated with Utt-B, while a drastic increase in the levels of these liver enzymes was observed in the control mice **(Fig. 6D-E)**. The renal function parameters such as levels of blood urea nitrogen and creatinine were much lower in the Utt-B-treated mice as compared to that of the control mice **(Supp. Fig. 1H-I)**. These results suggest that Utt-B stabilizes the hepatic and renal function in mice bearing metastatic HCC tumors. Further, immunohistochemical analysis of the tumor tissues of mice from the control and Utt-B groups revealed a significant reduction in the expression of MMPs, 2 and 9, the molecular markers associated with EMT, in the metastatic tumor tissues, suggesting the therapeutic role of Utt-B in preventing EMT and impeding the metastatic progression of HCC **(Fig. 6F)**. Immunohistochemical analysis demonstrates the elevated expression of p-EGFR, p-ERK1/2 and p-STAT3 in the control tumor tissues. However, there was a marked reduction in the expression of p-EGFR, p-ERK1/2 and p-STAT3 in the tumor tissue collected from the only tumor-bearing animal in the Utt-B group, compared to the control group mice, attesting the possible role of EGFR/ERK axis and STAT3 in regulating the robust anti-metastatic potential of Utt-B **(Fig. 6G)**.

**Figure 6:**
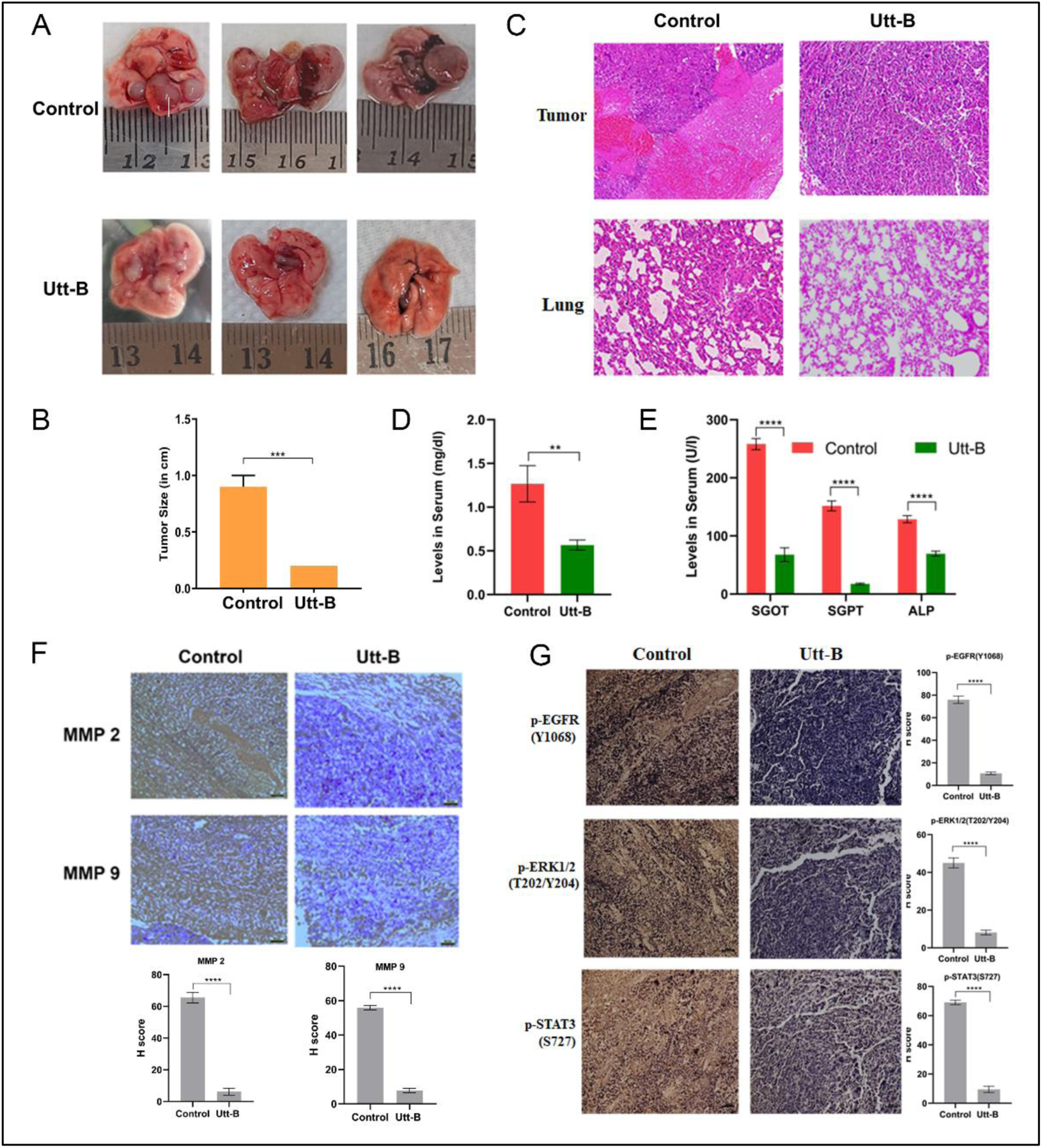
Utt-B mitigates the pulmonary metastasis of HCC, *in vivo,* via down-regulation of the EGFR/ERK axis and STAT3. **(A)** Representative images of lungs bearing metastatic tumors in control and Utt-B-treated mice from the murine metastasis model of HCC. **(B)** Graphical representation of the difference in the size of tumor nodules in mice from control and treatment groups. Student’s t test followed by Mann-Whitney test was used for statistical comparison between different groups. ***P ≤ 0.001. **(C)** Histopathological analysis of the metastatic tumor and corresponding lung tissues of control and Utt-B-treated mice from the murine metastasis model. **(D-E)** Levels of bilirubin and liver enzymes of animals from the control and Utt-B-treated groups as assessed by biochemical analysis of serum samples. Two-way ANOVA was used for statistical comparison between different groups. ****P ≤ 0.0001; **P ≤ 0.01. **(F)** Immunohistochemical analysis showing the relative tissue expression of EMT markers, MMP-2 and MMP-9 in the metastatic tumors of control and Utt-B-treated mice from the murine metastasis model of HCC. **(G)** Immunohistochemical analysis showing the relative tissue expression of p EGFR, p-ERK1/2 and p-STAT3 in the metastatic tumors of control and Utt-B-treated mice from Student’s t test analysis was used for statistical comparison between different groups. ****P ≤ 0.0001.

## Discussion

The transcriptomic and phosphoproteomic analyses in the current study have unraveled EGFR as a potential molecular target of Utt-B. A recent report by our team has documented that Utt-B gets encapsulated within the lipophilic layer of the extracellular vesicle (EV) membrane, forming complexes with cholesterol molecules [16]. Previous literature suggests that in the resting state, the EGFR organization on the plasma membrane is partly trapped in the cholesterol-containing domains of the plasma membrane [26]. The current *in silico* findings suggest that Utt-B exhibits a more stable binding with EGFR than mTOR. Therefore, it is possible that Utt-B binds to the extracellular domain of EGFR and induces the endocytosis of the receptor. Previous reports have demonstrated that endocytosis of EGFR impedes its activation by suppressing the autophosphorylation of its tyrosine kinase domains, in turn, leading to the inhibition of the signaling cascade occurring down-stream of EGFR [27]. Several studies have documented the therapeutic effects of EGFR tyrosine kinase inhibitors (TKIs) such as, gefitinib and erlotinib against HCC [28–30]. Kotzka *et al.,* have documented ERK and MAPK-mediated activation of SREBP-1 [8, 31]. The elevated expression of SREBP-1 is a characteristic feature of HCC and is responsible for excessive lipid biosynthesis and uptake, which exacerbate the development and progression of HCC [10]. Inhibition of SREBP-1 has yielded promising results in sensitizing HCC to radiofrequency ablation and sorafenib therapy [32, 33]. Our current findings suggest that Utt-B-mediated EGFR/ERK inhibition and the subsequent down-regulation of SREBP-1 could be one of the possible mechanisms underlying the exceptional anti-HCC potential of the compound. Clinical trials involving combinations of bevacizumab (anti-VEGF monoclonal antibody) with cetuximab (anti-EGFR antibody) [34], or erlotinib (EGFR TKI) [35] have augmented the therapeutic benefits of these agents against different types of cancer. We have previously reported the therapeutic efficacy and pharmacological safety of a novel combinatorial regimen involving Utt-B and sorafenib for combating HCC [36]. The simultaneous inhibition of EGFR and VEGF signaling by Utt-B and sorafenib, respectively, could be contributing to the additive therapeutic effect of this combination against HCC. A previous study suggests that the sustained up-regulation of EGFR and the concomitant STAT3 activation in HCC specimens correlates with the clinical stage of HCC [37]. Furthermore, the study has demonstrated that miR-491 prevents cancer stem cell-like properties and metastasis of HCC cells by blocking the EGFR-mediated activation of STAT3 [37]. A recent study has revealed the role of EMX1-FL in triggering metastasis via activation of EGFR/ERK signaling [38]. Another independent study has demonstrated that afatinib, an EGFR inhibitor, significantly inhibits EMT and tumorigenesis of Huh-7 cells via down-regulation of ERK/VEGF/MMP9 signaling pathway [39]. The current findings demonstrate the potential role of Utt-B in preventing EMT and the pulmonary metastasis of HCC, possibly via down-regulation of EGFR/ERK axis and STAT3 signaling.

Taken together, the current study elucidates the pharmacodynamics of Utt-B, provide compelling evidence on the robust anti-metastatic potential of the compound and highlight the key role of EGFR/ERK signaling axis in orchestrating the anti-HCC and anti-metastatic effects of Utt-B **(Fig. 7).** However, gene over-expression and/or silencing studies are highly warranted to establish EGFR as the ‘master-regulator’ governing the anti-HCC and anti-metastatic potential of Utt-B. The findings of the present study attest to the remarkable therapeutic efficacy of Utt-B as a promising drug candidate against primary and metastatic HCC and highlight its potential to be evaluated in the clinics for the benefit of HCC patients having limited prognosis and therapeutic options.

**Figure 7:**
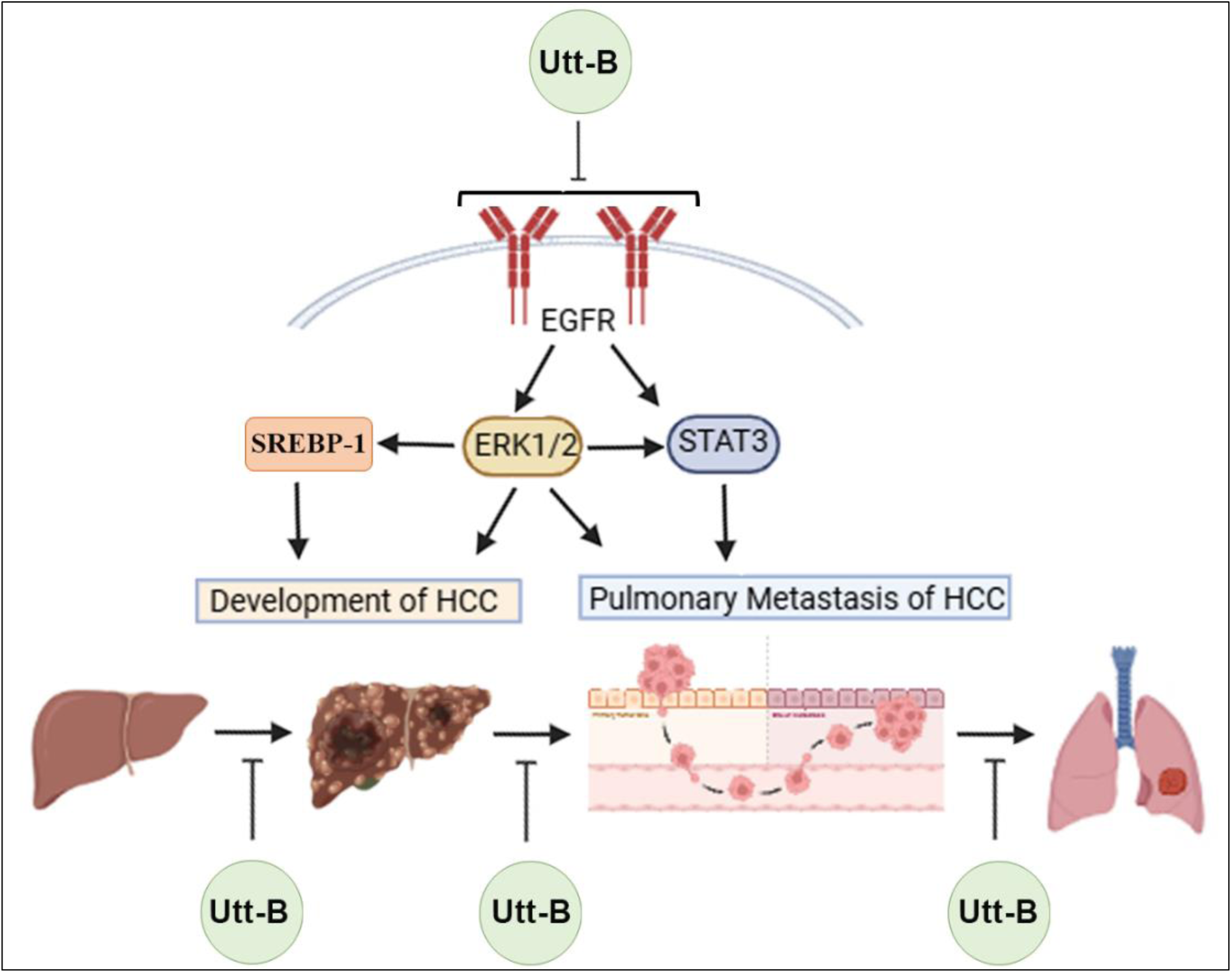
Uttroside B, a US FDA-designated ‘Orphan Drug’ against HCC mitigates the development and pulmonary metastasis of HCC via the down-regulation of EGFR/ERK axis.

## Ethical Declaration

### Conflict of Interest

The authors declare no competing interest.

### Animal Ethical Clearance

Studies involving experiments with animals were conducted in accordance with ARRIVE guidelines and institutional guidelines under the approval from Institutional Animal Ethics Committee, Rajiv Gandhi Centre for Biotechnology. (CPCSEA Number: 326/GO/ReBiBt/S/2001/CPCSEA). The animal experiments carried out as part of this study has been approved by the Institutional Animal Ethics Committee, IAEC No: IAEC/765/RUBY/2019 and IAEC/944/HARI/2023.

### Funding

This work was primarily funded by the Intramural Research Funding provided by the Department of Biotechnology, Government of India to RGCB. The authors also acknowledge the financial support received from DST-SERB to Dr. Ruby John Anto and Dr. K B Harikumar, and DBT-MK BHAN Young Researcher fellowship for the *in vitro* studies on the mechanism of Utt-B, for the Young Researcher fellowship awarded to Dr. Kalishwaralal K and Junior/Senior Research Fellowship to Dr. Keerthana CK. The funders had no role in designing the current study.

### Data Availability

Raw data compliant with the institutional confidentiality policies are available upon request. Data requests should be sent to the corresponding author.

## Acknowledgement

We thank Dr. Archana S, Dr. Arya Aravind, Dr. Queenie for the immense support provided in carrying out the animal experiments. We also acknowledge the help provided by the staff at the Animal Research facility and Core Research facilities of RGCB for the successful completion of all the experiments and the service provider Theracues for the completion of Nanostring experiment. The Figure 7 included in this manuscript has been created using BioRender.

## Author contribution

Concept and Study design: RJA; Acquisition of data: CKK, SS, AN, TPR, NHH, MS, KK, SUA, LRN, SSV, AV, JS, SJ, AJS, AP, VSJ; Formal Analysis and interpretation of the data: RJA, SS, TKM, SC, RSL; Writing the manuscript: CKK, RJA; Funding acquisition: RJA, KBH, KK; Review and editing: NPA, NI, RSL, KBH and RJA. All the authors have read and agreed to the current version of the manuscript.

---

**Supplementary Table 1:**
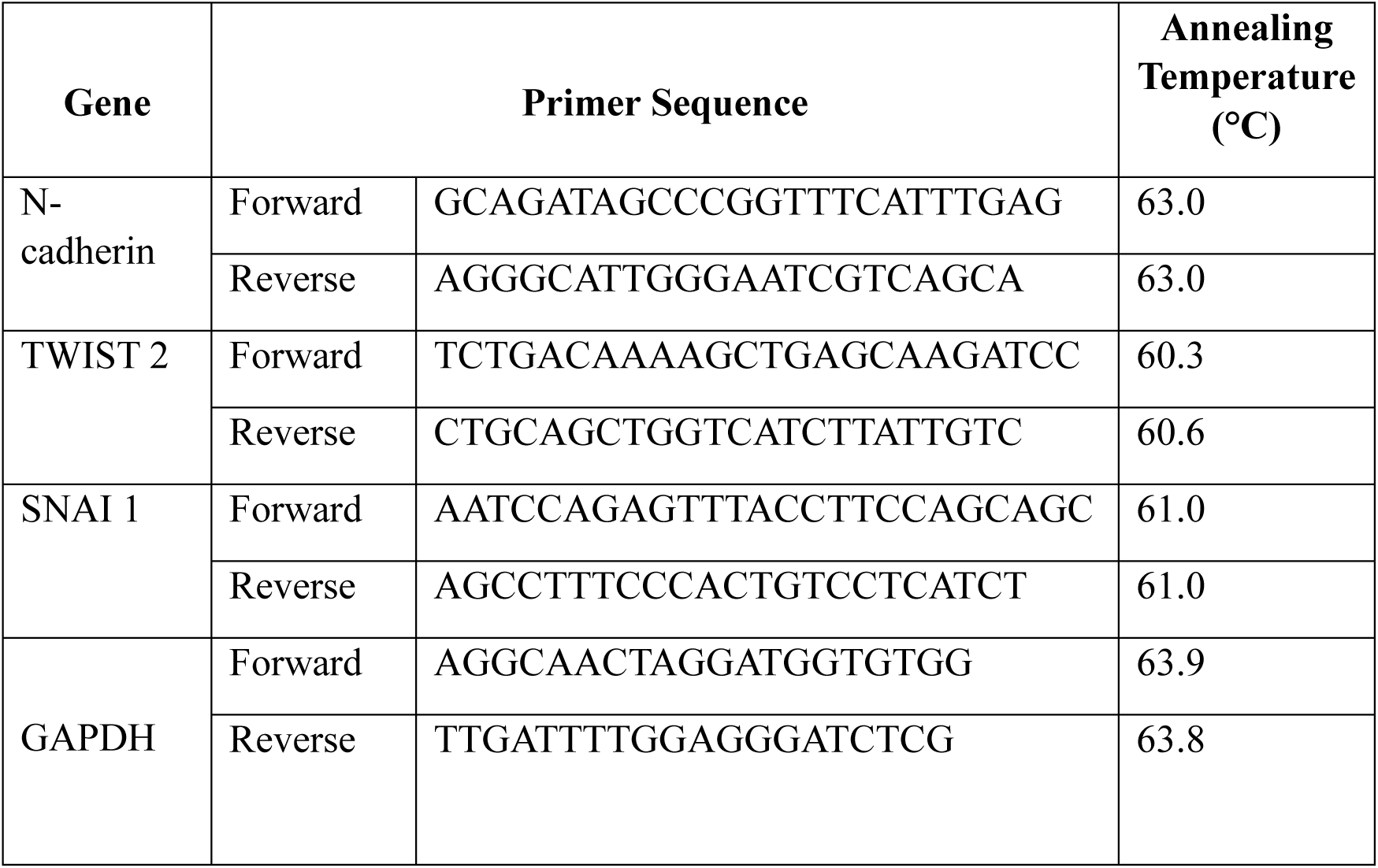
Details of human primers used in the study.

**Supplementary Figure 1:**
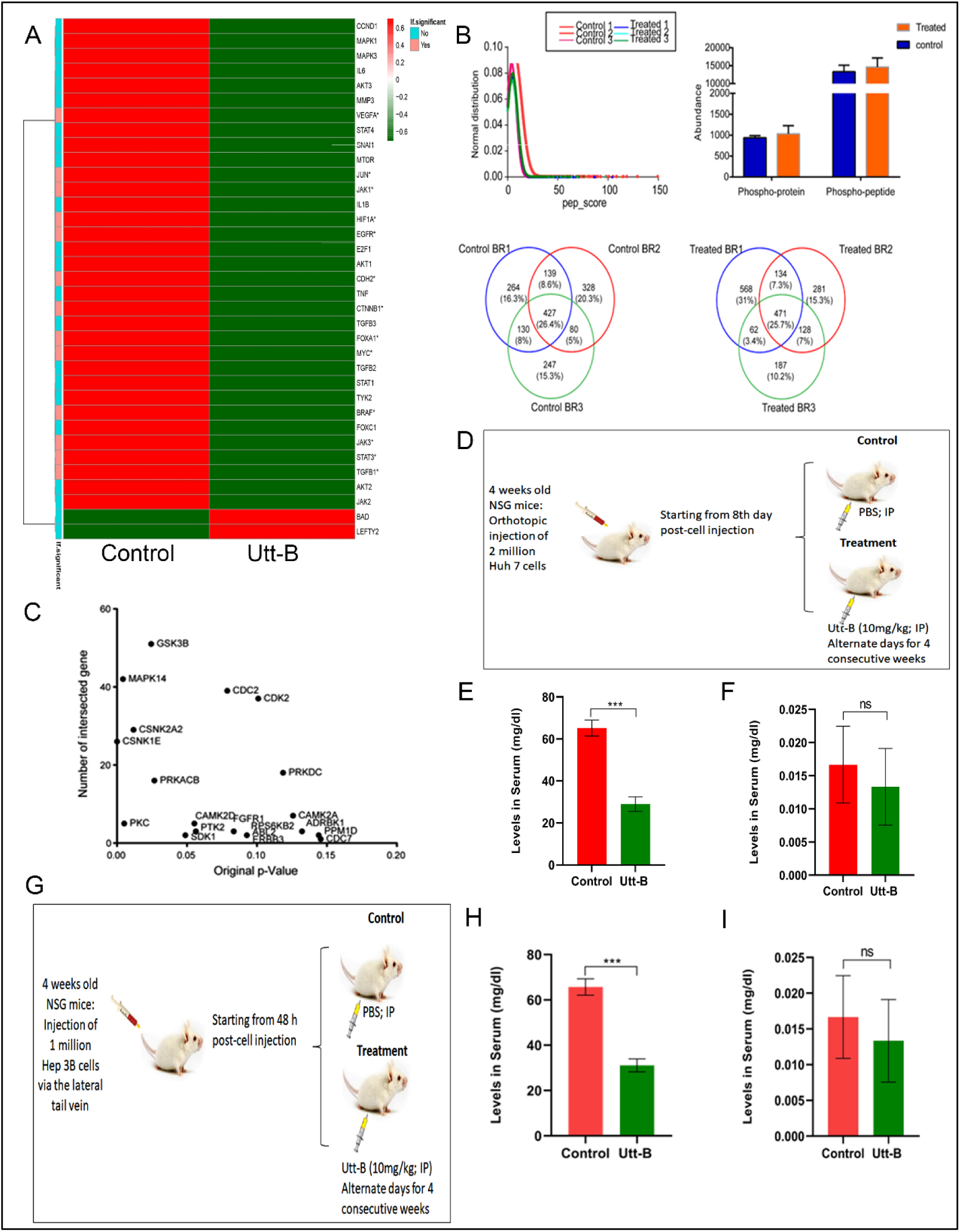
Utt-B down-regulates major survival pathways in HepG2 cells and stabilizes the renal function in murine models of primary and metastatic HCC. **(A)** Representative heat map of the differentially regulated genes in Utt-B-treated HepG2 cells with respect to the control and graph showing commonly enriched kinases in both control and Utt-B groups. **(B)** QC data of phosphoproteomics analysis and data depicting the number of differentially expressed phosphoproteins in the control and Utt-B-treated HepG2 cells in three biological replicates. **(C)** Kinome extraction data showing the enriched kinases common to both control and Utt-B treatment groups. **(D)** Schematic representation of the experimental design for the orthotopic xenograft model of HCC, *in vivo*. **(E-F)** Renal function profile of animals from the control and Utt-B treated groups from the orthotopic xenograft model of HCC, as assessed by biochemical analysis of serum samples. **(G)** Schematic representation of the experimental design for the murine metastasis model of HCC, *in vivo*. **(H-I)** Renal function profile of animals from the control and Utt-B-treated groups from the murine metastasis model of HCC, as assessed by biochemical analysis of serum samples. Student’s t test analysis was used for statistical comparison between different groups. ***P ≤ 0.001; ns-non-significant.

## Notes

### Competing Interest Statement

The authors have declared no competing interest.

